# Reciprocal regulatory interactions between class I TCP transcription factors and ABA signaling balance growth and stress responses in Arabidopsis

**DOI:** 10.64898/2026.06.17.732975

**Authors:** Alejo Canello, Juana Stipech, Antonela L. Alem, Martina N. Fus, Daniel H. Gonzalez, Ivana L. Viola

**Affiliations:** Instituto de Agrobiotecnología del Litoral (CONICET-UNL), Cátedra de Biología Celular y Molecular, Facultad de Bioquímica y Ciencias Biológicas, Universidad Nacional del Litoral, 3000 Santa Fe, Argentina

**Keywords:** *Arabidopsis thaliana*, TCP15, ABI5, AFP2, ABA signaling, salt stress responses

## Abstract

Abscisic acid (ABA) plays a critical role in regulating plant responses to abiotic stress by modulating various physiological processes through complex molecular networks. TEOSINTE BRANCHED1/CYCLOIDEA/PROLIFERATING CELL FACTORS (TCP) transcription factors are known developmental regulators, but their roles in ABA signaling and abiotic stress responses remain poorly understood. We addressed this question by analyzing the closely related *Arabidopsis thaliana* class I TCPs TCP14 and TCP15 using a variety of genetic and molecular approaches. We found that TCP14 and TCP15 negatively influence ABA and salt stress responses. They function by directly activating genes encoding negative regulators of ABA signaling, such as *ABA-INSENSITIVE FIVE-BINDING PROTEIN 2* (*AFP2*), thereby suppressing the expression of *ABA-INSENSITIVE 5* (*ABI5*) and downstream ABA-responsive genes to inhibit ABA responses under non-stressful conditions. Notably, the TCPs are targets of ABA-mediated regulation, as ABA negatively affects TCP14 and TCP15 protein abundance, providing a mechanism to limit TCP-dependent transcriptional growth responses and de-repress ABA-signaling pathways during times of stress. We propose that the antagonistic interplay between class I TCPs and ABA may serve to fine-tune plant growth and stress responses according to environmental conditions, positioning TCP14 and TCP15 as crucial players in balancing plant developmental progression and stress adaptation.

## 1. Introduction

Plants, as sessile organisms, are continually exposed to environmental fluctuations and various abiotic stresses, including drought, salinity and extreme temperatures, which can significantly affect growth, development, and productivity. To mitigate these effects, plants have evolved complex signaling networks that integrate environmental cues with physiological and developmental responses. Among these, the phytohormone abscisic acid (ABA) is a central regulator, orchestrating adaptive responses such as stomatal closure, osmotic adjustment, and transcriptional reprogramming of stress-related genes (Koornneef *et al*., 1984; Finkelstein & Lynch, 2000; Tuteja, 2007). Under optimal growth conditions, ABA levels remain relatively low in most plant tissues. However, abiotic stresses, as drought and salinity, rapidly induce ABA biosynthesis through the activation of 9-cis-epoxycarotenoid dioxygenase (*NCED*) genes, leading to ABA accumulation. Elevated ABA levels are sensed by the PYRABACTIN RESISTANCE1 (PYR1)/PYR1-LIKE (PYL)/REGULATORY COMPONENTS OF ABA RECEPTORS (RCAR) receptors, which inhibit clade A protein phosphatase 2Cs (PP2Cs), thereby activating SNF1-RELATED PROTEIN KINASE2 (SnRK2) family kinases (Ma *et al*., 2009; Park *et al*., 2009; Umezawa *et al*., 2009; Fujii *et al*., 2009). These kinases phosphorylate and activate ABA-responsive transcription factors, thereby controlling the expression of many downstream ABA-responsive genes (Kobayashi *et al*., 2005; Furihata *et al*., 2006; Fujii *et al*., 2007; Nakashima *et al*., 2009).

Genetic studies have identified numerous regulators of ABA signaling in Arabidopsis. Screens for ABA-insensitive (ABI) mutants revealed key regulators, such as *ABI3*, *ABI4*, and *ABI5*, which encode transcription factors from the B3, APETALA2/ERF, and basic leucine zipper (bZIP) families, respectively. These transcription factors positively regulate ABA signaling and response, facilitating growth arrest under unfavorable conditions (Giraudat *et al*., 1992; Finkelstein & Lynch, 2000; Finkelstein *et al*., 2002; Skubacz *et al*., 2016; Chandrasekaran *et al*., 2020). For instance, ABI5 regulates many ABA-dependent physiological responses, as seed dormancy, germination, cotyledon greening, root development, and leaf senescence, by binding to G-box-type ABA response elements (ABREs) in target genes and also to its own promoter (Choi *et al*., 2000; Finkelstein & Lynch, 2000; Lopez-Molina *et al*., 2001; Carles *et al*., 2002; Nakashima *et al*., 2006). Meanwhile, other factors negatively modulate ABA signaling at the transcriptional and post-transcriptional level. For example, the clade A PP2Cs ABI1 and ABI2 dephosphorylate and deactivate SnRK2 kinases (Leung *et al*., 1997), and ABI4 and ABI5 protein stability is affected via ubiquitination and sumoylation (Miura *et al*., 2009; Liu & Stone, 2013; Du *et al*., 2025), then attenuating ABA responses. In addition, different transcription factors act at the transcriptional level to inhibit ABA signaling. The WRKY transcription factors WRKY40, WRKY18, and WRKY60 repress the expression of a set of ABA-responsive and signaling genes, such as *ABI5*, during seed germination and seedling growth under non-stress conditions (Shang *et al*., 2010; Wang *et al*., 2021). Similarly, the BBX protein BBX21 interferes with ABI5 self-activation (Xu *et al*., 2014; Kang *et al*., 2018) and the MADS-box factor AGL16 negatively regulates the expression of certain salt stress-responsive and ABA-signaling genes (Zhao *et al*., 2020). In addition, the AP2/ERF transcription factor RELATED TO ABI3/VP1 (RAV1) plays important roles in ABA signaling by directly repressing *ABI3*, *ABI4*, and *ABI5* expression during seed germination and early seedling development (Feng *et al*., 2014). Furthermore, ABA response is inhibited by a small family of ABA-insensitive five binding proteins (AFPs), such as AFP1 and AFP2, which interact with ABI5 promoting its degradation and reducing ABA sensitivity during seed germination and early seedling development (Lopez-Molina *et al*., 2001, 2003; Garcia *et al*., 2008).

TEOSINTE BRANCHED1/CYCLOIDEA/PCF (TCP) is a plant-exclusive transcription factor family that plays essential roles in regulating a wide range of developmental processes (Cubas *et al*., 1999; Uberti Manassero *et al*., 2013; Viola *et al*., 2023). The family is defined by a conserved domain involved in DNA binding and dimerization that adopts a unique structure of three consecutive short β-strands rich in basic amino acids, followed by a helix-loop-helix structure (Sun *et al*., 2020). Based on sequence similarity, TCPs are classified into two main classes: I and II, comprising 11 and 13 members in Arabidopsis, respectively (Cubas *et al*., 1999). Despite structural similarities, TCPs exhibit both redundant and specialized functions. Moreover, opposite effects have been reported in certain processes, as cell cycle control and flowering, which would explain the maintenance of multiple members of this family across species (Viola *et al*., 2023; Viola & Gonzalez, 2023). Initially characterized for their roles in processes such as cell proliferation and leaf morphogenesis (Sarvepalli & Nath, 2018), accumulated evidence indicates that TCPs are important modulators of hormonal signaling pathways, including auxins, gibberellins, and cytokinins (Nicolas & Cubas, 2016; Danisman, 2016; Viola *et al*., 2023), and recent reports indicate that certain TCPs may also be involved in ABA-related pathways. For instance, the Arabidopsis class II TCP TCP13 regulates plant growth in leaves and roots to confer tolerance under dehydration stress conditions.

Mutants in *TCP13* show ABA-insensitive root growth and reduced dehydration-inducible gene expression, whereas TCP13 overexpression decreases water loss from leaves enhancing dehydration tolerance (Urano *et al*., 2022). In addition, class I TCPs from several species, such as PeTCP10, VuTCP9, OsTCP19 and OsPCF2, were shown to enhance stress tolerance and/or ABA sensitivity when overexpressed in Arabidopsis (Mukhopadhyay *et al*., 2015; Almeida *et al*., 2017; Liu *et al*., 2020; Mishra *et al*., 2021; Xu *et al*., 2021), whereas a negative role in ABA biosynthesis or signaling has been observed for GhTCP19, StTCP15 and MdTCP46 in gladiolus, potato and apple, respectively (Wu *et al*., 2019; Wang *et al*., 2022; Liu *et al*., 2022). These results suggest that different members of the TCP family play distinct roles in ABA responses. However, despite these findings, the molecular mechanisms through which TCP members integrate into ABA signaling and abiotic stress responses remain mostly unknown.

TCP14 and TCP15 are two closely related class I TCPs from Arabidopsis that redundantly participate in the control of diverse developmental processes, as germination, seedling establishment, flowering, aerial epidermis and flower development, and branching (Tatematsu *et al*., 2008; Resentini *et al*., 2015; Lucero *et al*., 2017; Zhang *et al*., 2019; Camoirano *et al*., 2020; Alem *et al*., 2022; Gastaldi *et al*., 2023, 2024). They act either as activators by binding to TCP-binding sites in target genes or by interacting with other proteins to modulate gene expression and downstream signaling pathways. In addition, these class I TCPs modulate hormonal pathways, as those of auxins and gibberellins, and responses to environmental conditions, as illumination during de-etiolation, high light intensity and increased temperature (Uberti-Manassero *et al*., 2012; Davière *et al*., 2014; Resentini *et al*., 2015; Viola *et al*., 2016; Ferrero *et al*., 2019, 2021; Gastaldi *et al*., 2020; Alem *et al*., 2022). It was also shown that they are important regulators in effector-triggered immunity (Kim *et al*., 2014; Li *et al*., 2018; Zhang *et al*., 2018; Ceulemans *et al*., 2021). In this study, we found that TCP14 and TCP15 are negative regulators of ABA signaling and salt stress responses in Arabidopsis. We show that these transcription factors suppress the expression of several ABA-responsive genes under non-stress conditions, acting upstream of *ABI5* and inducing the expression of negative regulators of ABA signaling, such as *AFP2*, *RAV1*, and *WRKY40*. We also found that ABA negatively impacts TCPs protein accumulation, which contributes to downregulation of the expression of TCP target genes involved in plant growth and the de-repression of ABA signaling under stress conditions. Altogether, our findings reveal reciprocal regulatory mechanisms between class I TCPs and ABA that ensure a proper balance between plant growth and stress responses in Arabidopsis, contributing to its developmental plasticity under fluctuating environmental conditions.

## 2. Materials and Methods

### 2.1. Plant material

All experiments were performed on Arabidopsis thaliana accession Col-0. Mutant lines tcp14-4, tcp14-6, tcp15-1, tcp15-3, tcp14-4 tcp15-3, abi5-8, the pTCP14::TCP14-GUS, pTCP15::TCP15-GUS-GFP, 35S::TCP15-RFP and 35S::YFP-AFP2 lines as well as plants that express estradiol inducible TCP15-GFP driven under the 35SCaMV promoter in the tcp15-3 background, 35S::TCP15-GFPi tcp15-3, were previously described (Kieffer et al., 2011; Zheng et al., 2012; Viola et al., 2016; Lynch et al., 2017; Ferrero et al., 2021; Alem et al., 2022). tcp14-4 tcp15-3 abi5-8 and 35S::YFP-AFP2 tcp14-4 tcp15-3 lines were generated by crossing. Primers used for genotyping are listed in Table S1.

### 2.2. Plant growth conditions and treatments

Plants were grown on soil or in plates containing 0.5X Murashige and Skoog (MS) medium and 0.8% agar at 23°C under long-day conditions (16 h light/8 h dark) at a light intensity of 100 µmol m-2 s-1. All seeds were surface-sterilized and stratified at 4°C for 4 days in the dark to synchronize germination. For tests of the ABA and NaCl effect on germination and seedling establishment, stratified seeds were grown on one-half-strength MS medium with or without ABA or NaCl for the times indicated in the Figures. For tests root elongation under ABA or NaCl treatments, 3-d-old seedlings vertically grown were transferred to MS medium supplemented with different concentrations of ABA or NaCl. Primary root length was measured 5d after transfer. For tests hypocotyl length and cotyledon expansion under ABA, seedlings were grown as described in the Figures. Measurements were performed as described in Ferrero *et al*. (2019) and Alem *et al*. (2022, 2025), respectively. For growth under non-lethal NaCl concentrations in soil, pots were soaked in water (control) or 60 mM NaCl before planting. This watering condition was maintained for 12 d. Then, NaCl concentration was increased to 100 mM until the end of the experiment. For high-NaCl treatments, plants were grown for 12 d under control conditions. Twelve days after sowing, subirrigation was carried out with a 50 mM NaCl solution, followed by weekly subirrigations with increasing NaCl concentrations (100, 150, and 200 mM) until the end of the experiment. ABA (Sigma) and NaCl (Merck) stock solutions were prepared according to the manufacturer’s recommendation.

### 2.3. RNA isolation and RT-qPCR analysis

RNA extractions from seedlings were performed as described (Chang *et al*., 1993). Reverse transcription (RT) was performed with an oligodTv primer and MMLV reverse transcriptase (PBL, Argentina) on 1.5–2.0 µg of total RNA. qPCR was performed in a StepOnePlus^TM^ Real Time PCR System thermocycler (LifeTechnologies) using the specific primers listed in Table S1 and SYBR Green detection. A comparative Ct method and three biological replicates were used to calculate relative transcript levels, with *ACT2* and *ACT8* actin genes as normalizers (Charrier *et al*., 2002).

### 2.4. Chromatin immunoprecipitation

Chromatin immunoprecipitation (ChIP) assays were performed as described in Alem *et al*. (2022) with anti-GFP (Abcam ab6556) or anti-IgG (Abcam ab6702) antibodies on chromatin prepared from 6-day-old *35S::TCP15-GFPi tcp15-3* seedlings grown under long-day conditions in liquid medium. TCP15 expression was induced by treating 5-d-old seedlings with 50 µM estradiol for 24 h. After cross-linking and extraction, chromatin was sonicated in a Bioruptor Pico water bath (Diagenode; 30s on/30s off pulses, at high intensity for 10 cycles, using Bioruptor microtubes). For immunoprecipitation, samples were incubated for 12 h at 4°C with Protein A Dynabeads (Invitrogen) pre-coated with the corresponding antibodies. Immunoprecipitated DNA was recovered using a phenol:chloroform:isoamyl alcohol mix (25:24:1) followed by ethanol precipitation. Untreated sonicated chromatin was processed in parallel and considered the input sample. DNA was analyzed by qPCR using primers listed in Table S1 and the SsoAdvanced Universal SYBR Green Supermix (Bio-Rad). The results presented are from three biological replicates. Between-experiment variation due to differences in absolute enrichment values for all regions analyzed was removed by factor correction (Ruijter *et al*., 2015).

### 2.5. β-Glucuronidase assay

β-Glucuronidase (GUS) activity was analyzed by histochemical staining using the chromogenic substrate 5-bromo-4-chloro-3-indolyl-β-D-glucuronic acid (X-gluc) as described by (Hull & Devic, 1995). Whole seedlings were immersed in a 0.5 g L-1 X-gluc solution in 50 mM citrate-HCl buffer, pH 7.0, and 0.05% Triton X-100. Vacuum was applied for 5 min and reactions were incubated in darkness at 37°C for 16 h. Chlorophyll was removed by incubating samples in 70% ethanol.

### 2.6. Western-blot analysis

Protein extracts for Western blot analysis were prepared as previously described (Ferrero *et al*., 2019), separated through SDS-PAGE, and then transferred to Hybond-ECL (GE Healthcare Life Sciences) membranes. Membranes were stained with Ponceau S and subsequently probed with antibodies against RFP (Living colors DsRed Polyclonal Antibody, Clontech; dilution of 1:1000) and developed with anti-rabbit immunoglobulin conjugated with horseradish peroxidase using the SuperSignal West Pico Chemiluminescent Substrate (ThermoFisher Scientific, Waltham, MA, USA), as previously described in *Viola et al.* (2016).

### 2.7. Fluorescence microscopy

GFP fluorescence in *pTCP15::TCP15-GUS-GFP* and *35S::TCP15-GFPi tcp15-3* seedlings was analyzed using a confocal laser scanning microscope (TCS SP8, Leica Microsystems, Mannheim, Germany) with excitation at 488 nm and detection at 493-522 nm. *pTCP15::TCP15-GUS-GFP* seedlings were grown during 6 d in liquid MS medium under normal conditions and then exposed for 24 h in the absence or presence of 10 μM ABA. *35S::TCP15-GFP_i_ tcp15-3* seedlings were grown in medium containing 50 µM β-estradiol for 6 days and then transferred to media lacking β-estradiol and with or without ABA for 24 h. Images were acquired in the sequential mode using Leica LCS software (LAS AF). For quantification of fluorescence intensity, approximately 200 nuclei from 10 independent images were measured for each treatment. Corrected total cell fluorescence intensity was calculated as: integrated density - (selected area of cell x mean fluorescence of background readings) using ImageJ (http://rsbweb.nih.gov/ij/).

### 2.8. Transcriptomics and functional genomics analyses

For the analysis of genes differentially expressed in *pTCP15:TCP15-EAR* and ABA-treated wild type plants processed microarray and RNA-sequencing data from Lucero *et al*. (2015) and Zhu *et al*. (2020) were retrieved from NCBI (GSE57744 and GSE135607, respectively). Gene ontology (GO) enrichment analysis was performed with PlantRegMap (Tian *et al*., 2019) and AgriGO (Tian *et al*., 2017). The GO terms with a P-value < 0.05 after Bonferroni correction were considered as significantly enriched GO terms.

## 3. Results

### 3.1. TCP14 and TCP15 are involved in the response to ABA

To explore the role of class I TCPs in ABA signaling in Arabidopsis, we analyzed single and double loss-of-function mutants in *TCP14* and *TCP15* for ABA-regulated growth responses. We first examined seedling establishment (i.e. the percentage of seedlings with green and expanded cotyledons) in wild-type and mutant plants grown in the absence or presence of ABA. Whereas no differences between the genotypes were observed in the absence of ABA (control condition), establishment of the *tcp14-4 tcp15-3* mutant was significantly more affected than that of wild-type plants in the presence of 0.5 µM ABA (Figure 1A). In addition, primary root elongation in the mutant was more significantly inhibited than in the wild-type when 3-day-old seedlings were transferred for 5 days to medium containing 15 µM ABA (Figure 1B), indicating that loss-of-function of *TCP14* and *TCP15* increases ABA sensitivity. Meanwhile, *tcp14* and *tcp15* single mutants did not display noticeable changes in ABA sensitivity both in seedling establishment and root growth assays (Figure S1A,B), suggesting that *TCP14* and *TCP15* function redundantly in ABA responses in Arabidopsis. Next, we analyzed ABA sensitivity in plants overexpressing TCP15 fused to the red fluorescent protein (RFP). In contrast to the *tcp14 tcp15* double mutant, *35S::TCP15-RFP* seedlings showed significantly higher establishment capacity when grown in the presence of 0.5 µM ABA and less sensitivity to ABA in root elongation assays in comparison with wild-type plants (Figure 1A,B). Altogether, the results suggest that TCP14 and TCP15 attenuate the response to ABA. Since TCP14 and TCP15 were shown to be involved in the promotion of seedling growth (Alem *et al*., 2022), we also analyzed hypocotyl elongation of dark-grown seedlings and cotyledon expansion during de-etiolation, two processes that have been reported to be inhibited by ABA (Hayashi *et al*., 2014; Lorrai *et al*., 2018; Martín *et al*., 2025) and are affected in *tcp14 tcp15* seedlings grown in the absence of the hormone (Figure 1C,D). Notably, hypocotyl growth in *tcp14 tcp15* etiolated seedlings was proportionally less affected by ABA treatment than in the wild type (Figure 1C). As a consequence, the growth difference between wild-type and mutant seedlings was progressively abolished in the presence of the hormone. Similar results were obtained when analyzing hypocotyl elongation and cotyledon expansion in seedlings grown under illumination in the absence and presence of ABA (Figure 1D,E). The results suggest that the reduced growth of *tcp14 tcp15* mutant seedlings under basal conditions may be partly due to activation of ABA responses.

**Figure 1.**
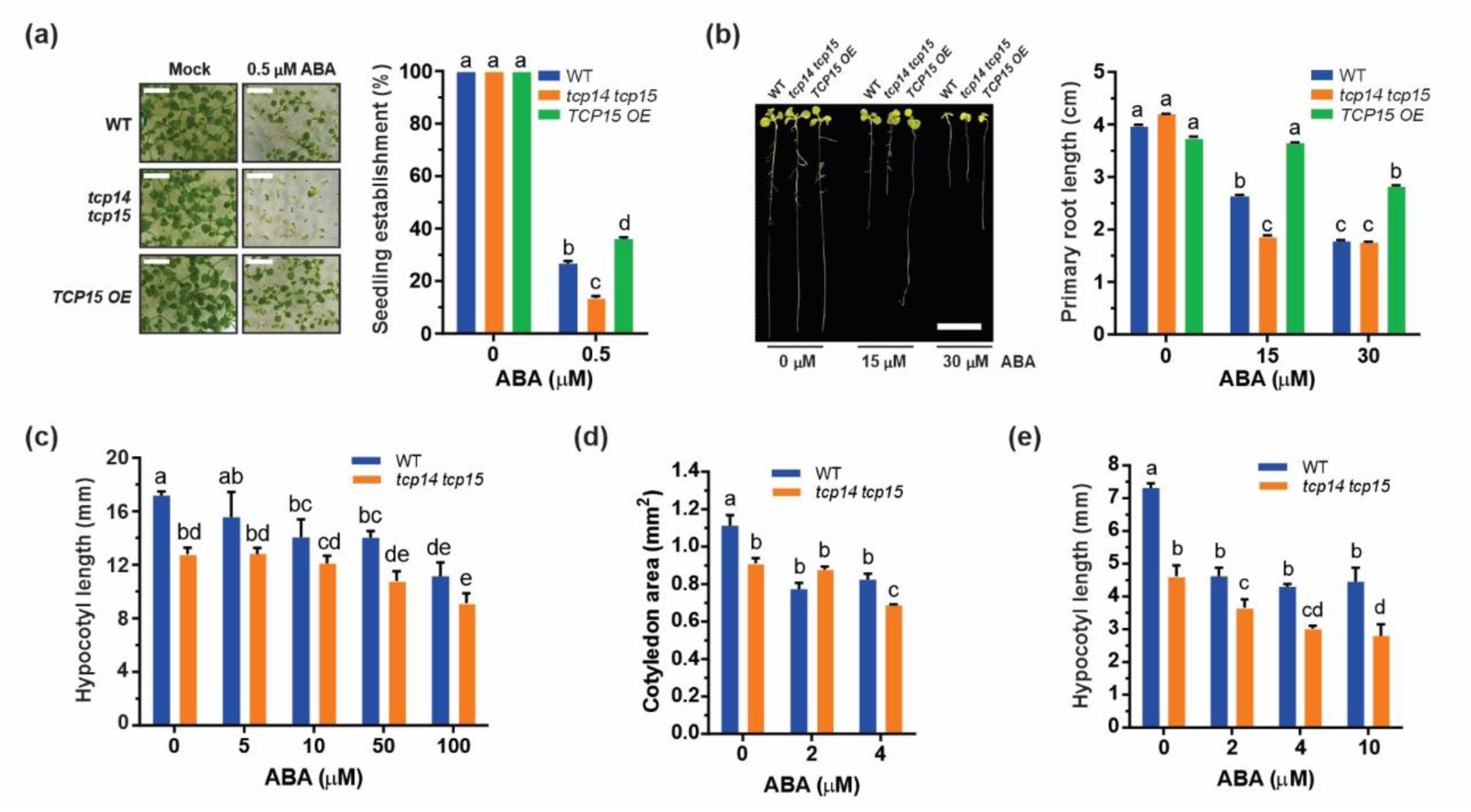
TCP14 and TCP15 participate in ABA responses. (A,B) Effect of exogenous ABA on seedling establishment (A) and primary root elongation (B) of wild-type (WT), *tcp14-4 tcp15-3* (*tcp14 tcp15*) and *35S::TCP15-RFP* (*TCP15 OE*) plants. In (A), seedlings were grown in the absence or presence of the indicated concentrations of ABA for 10 d. In (B), seedlings were grown for 3 d in the absence of ABA before being transferred to MS medium supplemented with the indicated concentrations of ABA. Root length was measured 5 d after transfer. Representative images are shown on the left of each panel (scale bars: 1 cm). (C) Hypocotyl length of WT and tcp14 tcp15 seedlings grown during 48 h in darkness and then transferred to MS medium supplemented with the indicated concentrations of ABA for 5 d under darkness. (D,E) Cotyledon area (d) and hypocotyl length (E) of the genotypes indicated in (C) grown during 48 h in darkness and then transferred to light in the presence of the indicated concentrations of ABA for 5 d. Bars show the mean±SD. Different letters denote statistically significant differences (P < 0.05; ANOVA). All the experiments were repeated three times with similar results.

### 3.2. Class I TCPs affect the expression of ABA-responsive genes

To further explore the role of class I TCPs in the ABA signaling pathway, we looked for genes related to this process in a global expression analysis of plants expressing a fusion of TCP15 to the EAR repressor motif under the control of the *TCP15* promoter (*pTCP15::TCP15-EAR*, Lucero *et al*., 2015). Gene Ontology (GO) analysis of differentially expressed genes revealed an enrichment in genes involved in ABA signaling, water deprivation and stress responses, as *ABI5*, *EARLY METHIONINE-LABELED6* (*EM6*) and *COLD-RESPONSIVE 6.6* (*COR6.6*/*KIN2*) (Table S2). Notably, most genes in these categories were upregulated in *pTCP15::TCP15-EAR* plants compared to wild-type (i.e. 81% for ABA-response genes, 77.5% for water-deprivation genes and 70% for stress-response genes) (Tables S3-S5), suggesting that ABA-related responses are induced in TCP15-EAR expressing plants. Next, we examined the expression of ABA-response genes in *tcp14 tcp15* mutant seedlings. We found that expression of the key positive regulators of ABA signaling *ABI4* and *ABI5* was significantly increased in *tcp14 tcp15* mutant seedlings compared to wild type (Figure 2A). In addition, the expression of the downstream ABA-responsive genes *EARLY METHIONINE-LABELED1* (*EM1*), *EM6*, *RESPONSIVE TO ABA18* (*RAB18*) and *RESPONSIVE TO DESICCATION29B* (*RD29B*) (Foster & Chua, 1999; Carles *et al*., 2002; Nakashima *et al*., 2006; Chen *et al*., 2014) was also significantly increased in *tcp14 tcp15* seedlings compared to wild type (Figure 2A), indicating that class I TCPs negatively modulate the expression of ABA responsive genes.

**Figure 2.**
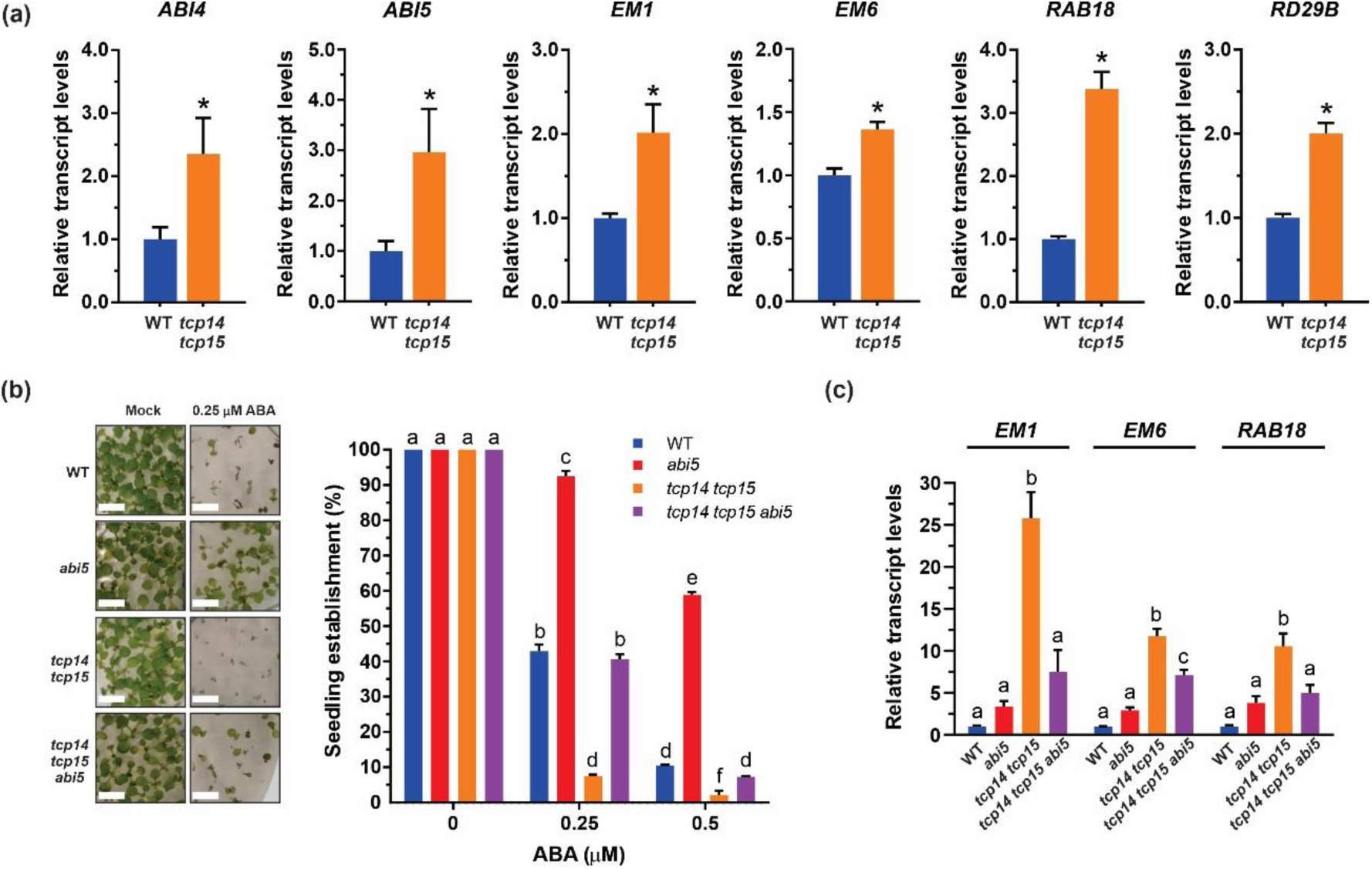
Class I TCPs modulate the expression of ABA-responsive gene and seedling sensitivity to ABA acting upstream of *ABI5*. (A) Transcript levels of the positive regulators of ABA signaling *ABI4* and *ABI5* and downstream ABA-responsive genes in 6-d-old wild-type (WT) and *tcp14-4 tcp15-3* (*tcp14 tcp15*) seedlings. Values are expressed relative to wild type. (B) Seedling establishment capacity of WT, *abi5-8* (*abi5*), *tcp14 tcp15* and *tcp14 tcp15 abi5* seedlings grown for 10 d in the absence or presence of the indicated concentrations of ABA. Representative images are shown on the left (scale bars: 1 cm). (C) Expression of ABI5-regulated genes in 2-d-old WT, *abi5*, *tcp14 tcp15* and *tcp14 tcp15 abi5* seedlings. Values are expressed relative to wild type. The bars indicate the mean±SE of three biological replicates. In (A), asterisks indicate significant differences compared with the wild type (P < 0.05; Student’s t-test). In (B,C), different letters denote significant differences (P < 0.05; ANOVA). All the experiments were repeated three times with similar results.

### 3.3. Class I TCPs function upstream of ABI5 in the ABA signaling pathway

Since ABI5 is a main factor controlling post-germination growth arrest and inhibition of seedling establishment in response to ABA, we hypothesized that the enhanced ABA sensitivity of *tcp14 tcp15* mutant seedlings could be partly due to the increased expression of *ABI5*. To test this, we generated a *tcp14-4 tcp15-3 abi5-8* triple mutant and evaluated establishment on MS medium supplemented with ABA. While the single *abi5* mutant and the double *tcp14 tcp15* mutant showed decreased and increased ABA sensitivity relative to wild type, respectively, the triple *tcp14 tcp15 abi5* mutant exhibited an ABA-sensitive phenotype similar to that of wild-type seedlings (Figure 2B). In addition, the observed increase in expression levels of the ABI5-regulated genes *EM1*, *EM6* and *RAB18* was mostly abolished after introducing the *abi5* mutation in the *tcp14 tcp15* background (Figure 2C), suggesting that *ABI5* acts downstream of class I TCPs. However, the fact that the *tcp14 tcp15 abi5* triple mutant did not exhibit the same ABA-insensitive phenotype of *abi5* mutant seedlings (Figure 2B) suggests that TCP14 and TCP15 also affect the ABA pathway through *ABI5*-independent mechanisms.

### 3.4. TCP15 promotes the expression of negative regulators of ABA signaling

Our results indicate that class I TCPs function as negative regulators of ABA signal transduction in Arabidopsis. However, as the transcriptional repressor form of TCP15 (TCP15-EAR) induces ABA-responsive genes, this regulation is likely indirect. In addition, most class I TCPs have been characterized as transcriptional activators (Viola *et al*., 2023), suggesting that they may act by promoting the expression of a repressor of the ABA pathway. To explore this, we sought for negative regulators of ABA responses whose mutation causes phenotypic and gene expression changes similar to those observed in *tcp14 tcp15* mutants and contain putative TCP binding sites (GGNCCC, Viola *et al*., 2011, 2012) in their promoters. Our analysis revealed that the transcription factor coding genes *ABA-INSENSITIVE FIVE-BINDING PROTEIN 2* (*AFP2*), *RELATED TO ABI3/VP1 1* (*RAV1*) and *WRKY DNA-BINDING PROTEIN 40* (*WRKY40*) meet these requirements. Interestingly, we found that the expression of these genes was significantly reduced in *tcp14 tcp15* mutant seedlings compared to the wild type, while it was significantly increased in plants overexpressing *TCP15* (Figure 3A). Then, we evaluated the binding of TCP15 to the promoters of these genes through a chromatin immunoprecipitation (ChIP) followed by qPCR assay using 6-day-old seedlings containing a β-estradiol-inducible construct for the expression of TCP15 fused to GFP (*35S::TCP15-GFP_i_*) and anti-GFP antibodies. We found that TCP15 associated to the regions containing TCP boxes in the *AFP2*, *RAV1* and *WRKY40* promoters (Figure 3B), indicating that these genes are direct TCP15 targets. This suggests that TCP15, and probably also TCP14, modulate ABA responses by inducing the expression of one or more negative regulators of the ABA signaling pathway. To provide further support for this, we chose AFP2 and analyzed whether *AFP2* overexpression could abolish the enhanced ABA sensitivity of the *tcp14 tcp15* mutant. We observed that introducing a construct for *AFP2* overexpression (*35S::YFP-AFP2*) restored seedling establishment of the *tcp14 tcp15* mutant under ABA treatment to wild-type levels (Figure 3C), suggesting that the decreased *AFP2* expression is partly responsible for the higher ABA sensitivity shown by *tcp14 tcp15* mutant plants. Nevertheless, *35S::YFP-AFP2 tcp14-4 tcp15-3* seedlings did not achieve the same level of ABA insensitivity of *35S::AFP2* seedlings (Figure 3C), indicating that other factors, probably *WRKY40* and *RAV1* among them, are also involved.

**Figure 3.**
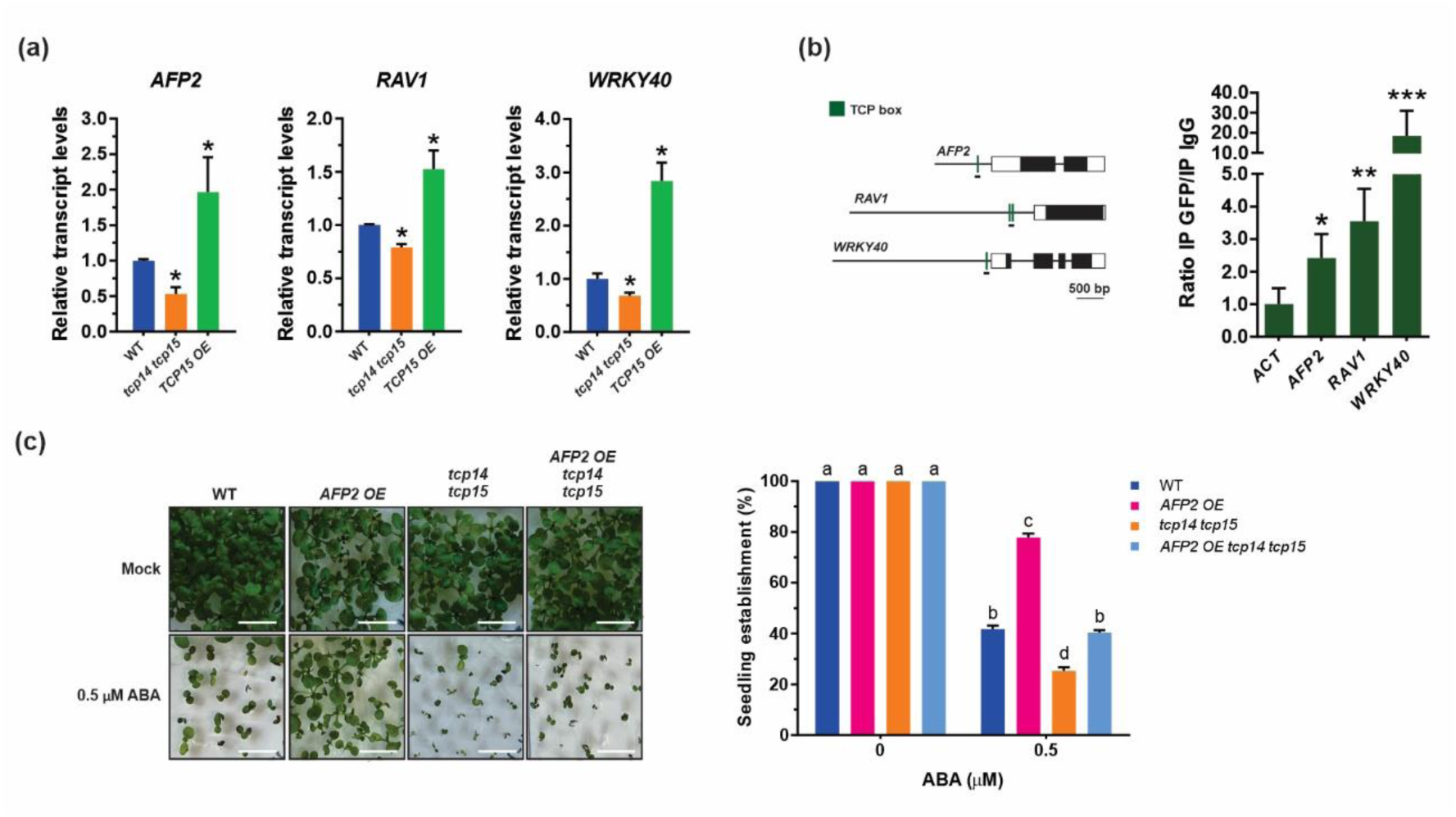
TCP15 induces the expression of negative regulators of ABA signaling. (A) Quantitative analysis of *AFP2*, *RAV1* and *WRKY40* transcript levels in wild-type (WT), *tcp14-4 tcp15-3* (*tcp14 tcp15*) and *35S::TCP15-RFP* (*TCP15 OE*) 6-d-old seedlings. Values are expressed relative to wild type. The bars indicate the mean±SE of three biological replicates. Asterisks indicate significant differences compared with the wild type (P < 0.05; Student’s t-test). (B) Association of TCP15 with the promoters of *AFP2*, *RAV1* and *WRKY40* analyzed by ChIP-qPCR. Left panel, diagram of the genes showing the positions of putative TEOSINTE BRANCHED1, CYCLOIDEA, and PCF (TCP) binding motifs (TCP box; GTGGGNCC, green rectangles) and the regions analyzed by ChIP-qPCR. Exons and untranslated transcribed regions are represented as black and white boxes, respectively. Right panel, ChIP-qPCR analysis of the binding of TCP15-GFP to the indicated promoters, performed in 6-d-old seedlings expressing an estradiol-inducible TCP15-GFP construct in *tcp15-3* background. A region of the *ACTIN2* gene (*ACT*) was used as negative control. Bars show the mean±SE of three biological replicates. Asterisks indicate significant differences compared with *ACTIN2* (P < 0.05; Student’s t-test). (C) Seedling establishment capacity of WT, *35S::YFP*-*AFP2* (*AFP2 OE*), *tcp14 tcp15* and *AFP2 OE tcp14 tcp15* seedlings grown in the absence or presence of 0.5 µM ABA. Different letters denote significant differences (P < 0.05; ANOVA). Representative images are shown on the left (scale bars: 1 cm). All the experiments were repeated three times with similar results.

Altogether, the results indicate that class I TCPs repress ABA signaling pathways in Arabidopsis by inducing the expression of negative regulators that impinge on the activity of *ABI5* and possibly other effectors of ABA responses. This mechanism would ensure a robust and coordinated suppression of ABA-responsive processes to prevent negative effects on growth in the absence of the hormone and avoid excessive responses to ABA under stress conditions.

### 3.5. Class I TCPs negatively regulate abiotic stress responses

At this point, we asked about the role of class I TCPs under physiological conditions that induce ABA signaling, as growth under excessive salt concentrations. We found that the *tcp14 tcp15* mutant displayed hypersensitivity to NaCl during seed germination (Figure 4A) and seedling establishment (Figure 4B), whereas *35S::TCP15-RFP* seedlings were less sensitive to salt than the wild-type (Figure 4A,B). In addition, when seeds germinated on MS medium for 3 days were transferred to medium supplemented with NaCl, the *tcp14 tcp15* mutant displayed stronger inhibition of primary root elongation in comparison with the wild type (Figure 4C), whereas *TCP15* overexpressing seedlings showed the opposite behavior (Figure 4D). The results are consistent with the phenotypes observed under ABA treatment, suggesting that the effect of NaCl is probably linked to an increase in ABA signaling during early seedling development.

**Figure 4.**
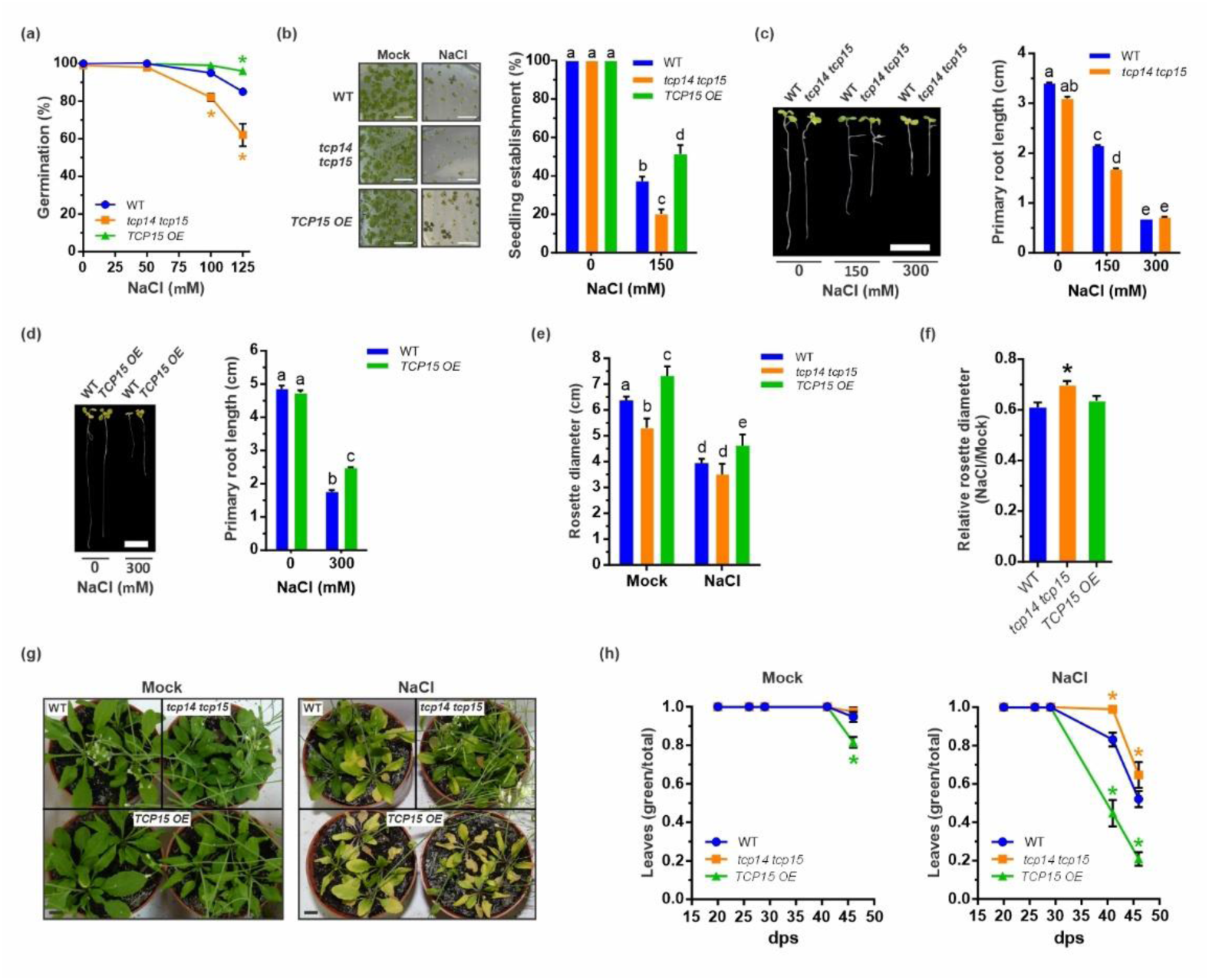
Class I TCPs negatively affect plant salt sensitivity. (A,B) Effect of NaCl on germination rate (A) and seedling establishment (B) after growth for 3 d (a) or 10 d (B) in the absence or the presence of the indicated concentrations of NaCl. In (B), representative images are shown on the left (scale bars: 1 cm). (C,D) Effect of NaCl on primary root elongation. Seedlings were grown for 3 d in the absence of NaCl and then transferred to MS medium supplemented with the indicated NaCl concentrations for 5 d. Representative images are shown on the left of each panel (scale bars: 1 cm). (E) Rosette diameter of plants grown for 24 d under control conditions (Mock) or under 100 mM NaCl. Bars show the mean ± SD (n=10). (F) NaCl sensitivity expressed as the rosette diameter under NaCl relative to control condition for the genotypes in (E). (G) Representative images of plants grown under control conditions (Mock) or exposed to increasing NaCl concentrations (50, 100, 150, 200 mM) from 10 d post-stratification (NaCl; see Materials and methods for details). Scale bars: 1 cm. (H) Ratio of green to total rosette leaves in plants grown as in (G). Measurements were made at the indicated time points. Values are the mean ± SD (n = 10). In (A,F,H), asterisks indicate significant differences compared with the wild type (P < 0.05; Student’s t-test). In (B-E), different letters denote significant differences (P < 0.05; ANOVA). All the experiments were repeated three times with similar results. WT, wild-type; *tcp14 tcp15*, *tcp14-4 tcp15-3*; *TCP15 OE*, *35S::TCP15-RFP*.

Next, we analyzed the response to high-salt treatments of rosette-stage plants. For this, plants were germinated and grown for 15 days in 60 mM NaCl and then irrigated with 100 mM NaCl as specified in Materials and Methods. Under control conditions (without NaCl), the *tcp14 tcp15* mutant displayed smaller rosettes compared to wild-type plants, while the opposite was observed for *TCP15* overexpressing plants (Figure 4E). The salt treatment significantly decreased the rosette diameter in all analyzed genotypes compared to control conditions (Figure 4E). However, the growth of *tcp14 tcp15* mutant plants was proportionally less inhibited by NaCl, reaching values similar to those of wild-type plants after the salt treatment (Figure 4E,F). Meanwhile, the relative effect of the salt treatment was similar to that observed for wild-type plants for *35S::TCP15-RFP* plants (Figure 4F). In addition, when plants were irrigated with a higher NaCl concentration (200 mM) after growth on soil for two weeks, *tcp14 tcp15* mutant plants exhibited greater tolerance than wild-type plants as judged by the proportion of leaves that stayed green after the treatment (Figure 4G,H). Meanwhile, *35S::TCP15-RFP* plants exhibited accelerated leaf yellowing, which was significantly more pronounced under NaCl treatment (Figure 4G,H). It can be speculated that the altered expression of ABA-responsive genes, which are critical for abiotic stress tolerance, would be responsible for the behavior of *tcp14 tcp15* and *TCP15* overexpressing plants under salt-stress inducing conditions. Altogether, the results suggest that the class I TCPs TCP14 and TCP15 function as negative regulators of responses to abiotic stress by downregulating the expression of key components of the ABA-response pathway.

### 3.6. ABA affects TCP protein accumulation at the post-transcriptional level

The fact that TCP14 and TCP15 negatively affect plant responses to stress poses the question of whether the activity of the TCPs is regulated by ABA and/or stress conditions to allow plants to display a proper response. To evaluate this, we first examined transcriptomic data from Arabidopsis plants exposed to ABA or high NaCl concentrations, and found no significant changes in the expression of *TCP14* or *TCP15* (Kuhn *et al*., 2008; Dinneny *et al*., 2008; Chan *et al*., 2011). Then, we analyzed *TCP14* and *TCP15* expression under ABA treatment by RT-qPCR. We found that *TCP14* transcript levels were decreased by ca. 40% when 6-day-old wild-type seedlings were treated with 10 µM ABA for 24 h (Figure 5A), suggesting that ABA represses *TCP14* expression in seedlings. However, *TCP15* transcript levels were not significantly affected by ABA (Figure 5A). To further explore this, we analyzed plants expressing the *TCP14* and *TCP15* coding regions fused to the *GUS* and *GFP-GUS* reporter genes, respectively, under the control of their native promoters. Analysis of *GUS* reporter activity revealed a strong decrease in *GUS* expression for both reporters after plants were treated with ABA or NaCl for 24 h (Figure 5B, S2). Since the TCP15 reporter construct also contains GFP, we analyzed GFP fluorescence in cotyledons and found that fluorescence intensity was also significantly reduced after ABA treatment (Figure 5C). Notably, the decrease in GUS and GFP levels caused by ABA seems to be significantly greater than the decrease in *TCP14* and *TCP15* transcript levels observed under the same conditions, suggesting the existence of regulation at the post-transcriptional level.

**Figure 5.**
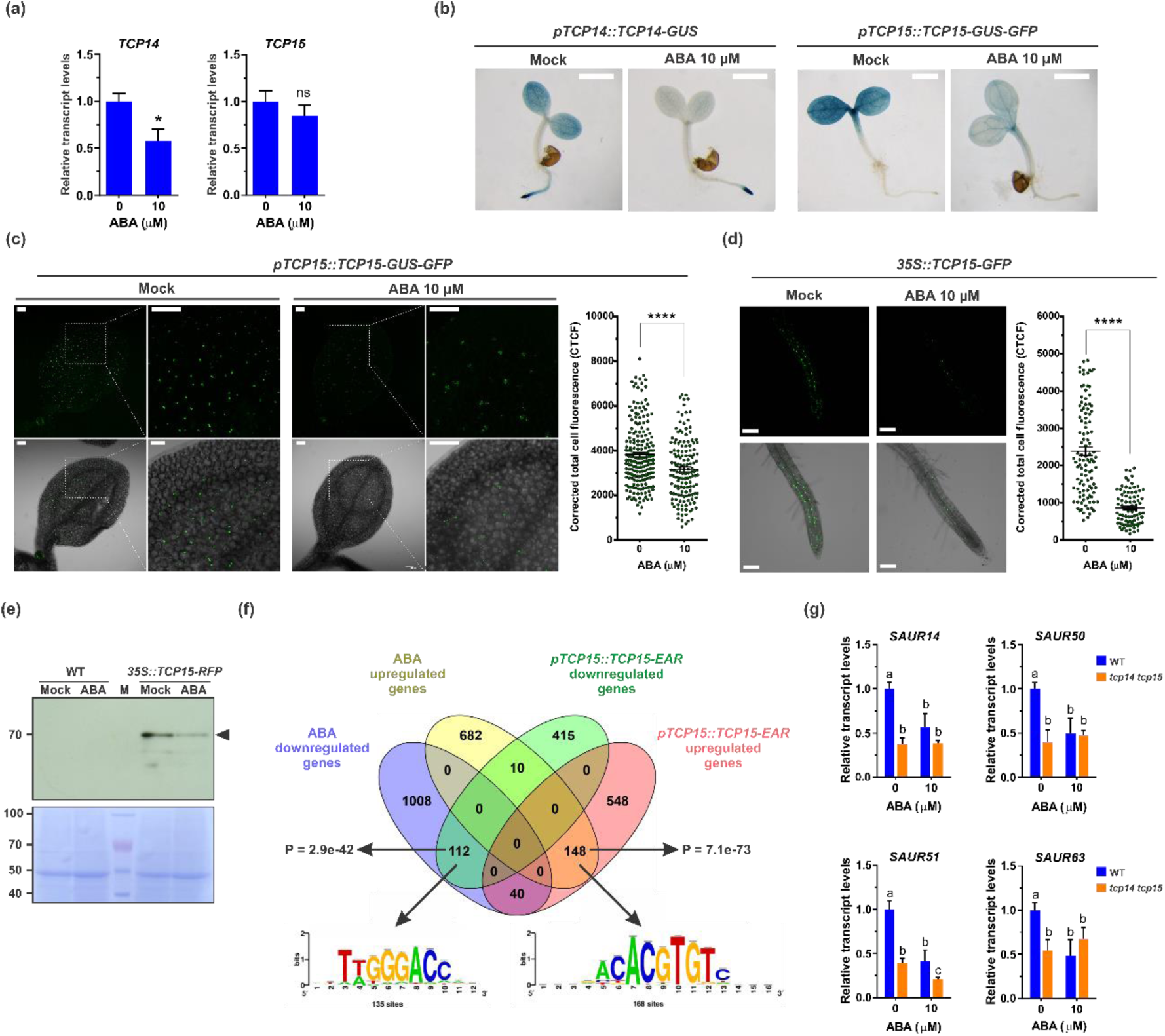
ABA downregulates TCP protein abundance and TCP15-target gene expression. (A) Quantitative analysis of *TCP14* and *TCP15* transcript levels in wild-type seedlings grown during 6 d in liquid MS medium under normal conditions and then incubated for 24 h in the absence or presence of 10 μM ABA. The bars indicate the mean±SE of three biological replicates. Asterisks indicate significant differences (P < 0.05; Student’s t-test). (B) TCP14 and TCP15 expression pattern analyzed by β-glucuronidase (GUS) histochemical staining of *pTCP14::TCP14-GUS* and *pTCP15::TCP15-GUS-GFP* seedlings grown as indicated in (a). Images are representative of 10 seedlings analyzed (scale bars: 1 mm). (C,D) Quantitative analysis of GFP fluorescence in seedlings harboring the *pTCP15::TCP15-GUS-GFP* (C) and the estradiol-inducible *35S::TCP15-GFP* construct (D) grown as described in the text. GFP fluorescence intensity was measured in ca. 200 nuclei from 10 independent images using the IMAGEJ software (http://rsb.info.nih.gov/ij/). Bars indicate the mean±SE and asterisks indicate significant differences (P < 0.05; Student’s t-test). Representative confocal microscopy images and zoom images used for fluorescence quantification are shown on the left, with lower panels showing merged images of GFP fluorescence and transmitted light (scale bars: 100 µm). (E) Western blot analysis with anti-RFP antibodies of TCP15-RFP protein levels in extracts obtained from wild type (WT, negative control) and *35S::TCP15-RFP* seedlings grown during 5 d in liquid MS medium under normal conditions and then treated with either 10 μM ABA (ABA) or DMSO (Mock) for 24 h, as indicated. Total proteins on the blotted membrane were stained with Coomassie blue after immunodetection (lower panel) as loading control. Numbers indicate molecular masses of molecular-weight standards. The black arrow indicates the expected migration of full-length TCP15-RFP. (F) Venn diagram of differentially expressed genes in plants expressing a repressor form of TCP15 under the control of the *TCP15* promoter (*pTCP15::TCP15-EAR*) and ABA-treated wild-type plants. P-values for the genes that are either downregulated or upregulated in both transcriptomes, calculated by a hypergeometric distribution test, are indicated. Logos of sequences enriched in the promoter regions of commonly downregulated (TGGGACC, class I TCP-binding motif) and upregulated (CACGTG, G-box/ABRE-like motif) genes are shown. Enriched sequences were analyzed using the oligoanalysis motif discovery tool from RSAT (Regulatory Sequence Analysis Tools; http://rsat.eead.csic.es/plants/) on sequences located 3-kb upstream of the transcription start site, preventing overlap with neighboring genes. (G) Quantitative analysis of growth-related TCP15-target gene transcript levels in WT and *tcp14-4 tcp15-3* seedlings grown as indicated in (e). The bars indicate the mean±SE of three biological replicates. Different letters denote significant differences (P < 0.05; ANOVA). All the experiments were repeated three times with similar results.

To further address possible post-transcriptional effects of ABA on TCP protein levels, we analyzed TCP15 accumulation in *35S::TCP15-GFP_i_* seedlings, expressing TCP15-GFP under the control of the *35SCaMV* constitutive promoter in a β-estradiol-inducible fashion. In this assay, *35S::TCP15-GFP_i_* seedlings were grown in medium containing 50 µM β-estradiol for 6 days and then transferred to media lacking β-estradiol and with or without ABA for 24 h. We observed a significant decrease in TCP15-GFP levels of ABA-treated seedlings compared to those kept under control conditions (Figure 5D). Additionally, proteins extracted from *35S::TCP15-RFP* seedlings grown under the same conditions were subjected to immunoblot analysis with anti-RFP antibodies. As shown in Figure 5E, ABA treatment also caused a decrease in TCP15-RFP protein levels in these plants. The results suggest that ABA post-transcriptionally affects the accumulation of the class I TCP proteins, probably acting on their stability.

To assess the impact of ABA-mediated TCP-downregulation on gene expression, we compared the transcriptomic profiles of wild-type plants treated with ABA (Zhu *et al*., 2020) and plants expressing a dominant repressor form of TCP15 (TCP15-EAR) (Lucero *et al*., 2015). We found significant overlap between the sets of genes whose expression is similarly affected in both conditions (Figure 5F). Gene Ontology (GO) analysis of genes commonly upregulated by ABA and TCP15-EAR revealed an enrichment in genes involved in several abiotic stresses and ABA responses (Table S6), reflecting the negative regulation of ABA-responsive genes exerted by TCP15. Interestingly, these genes showed an enrichment of G-box/ABRE-like motifs in their promoters, as expected for ABA-induced genes and the fact that TCP15 regulation of these genes is indirect (Figure 5F). Meanwhile, GO analysis of genes downregulated by both ABA and TCP15-EAR revealed an enrichment in growth-related terms, as "growth regulation", "organ growth", and "developmental growth" (Table S7). These sets are enriched in known TCP15-target genes, as *HBI1*, *PRE1*, *PRE6*, *SAURs* and *EXPANSINs* (Ferrero *et al*., 2019, 2021; Gastaldi *et al*., 2020; Alem *et al*., 2022) (Table S8). Moreover, the promoters of these genes show an enrichment of the class I TCP recognition motif TGGGACC (Viola *et al*., 2011, 2012). These observations suggest that ABA-mediated repression of several growth-promoting genes operates through class I TCPs. To validate this, we analyzed the expression of a group of *SAUR* genes that are directly induced by TCP15 and other TCPs to promote growth (Koyama *et al*., 2010, 2025; Dong *et al*., 2019; Gastaldi *et al*., 2020; Ferrero *et al*., 2021; Alem *et al*., 2022, 2025). We found that these TCP15 targets were significantly repressed by ABA in wild-type seedlings (Figure 5G). Meanwhile, *tcp14 tcp15* plants showed reduced *SAURs* gene expression, similar to that of ABA-treated wild-type seedlings, in the absence of ABA, and no further decrease after ABA treatment (Figure 5G), suggesting that ABA represses *SAURs* expression through TCP14 and TCP15. Altogether, the results indicate that ABA negatively affects TCP14 and TCP15 accumulation as a mechanism to promote ABA-dependent growth arrest and stress adaptation.

## 4. Discussion

### 4.1. TCP14 and TCP15 are negative regulators of ABA signaling in Arabidopsis

In this study, we explore the role of the class I TCP transcription factors TCP14 and TCP15 from Arabidopsis in ABA signaling. We found that a *tcp14 tcp15* double mutant shows enhanced sensitivity to ABA during seedling establishment and primary root elongation, whereas ectopic expression of TCP15 decreases ABA sensitivity, indicating a suppressive role for these TCPs on ABA responses. Interestingly, single mutants in *TCP14* and *TCP15* did not exhibit altered ABA sensitivity, suggesting that they function redundantly in ABA responses. Overlapping roles between TCP14 and TCP15 were previously observed in the biosynthesis and signaling of different hormones, as auxin, cytokinin and gibberellin, and under various developmental and environmental conditions, indicating that these transcription factors exert highly redundant roles in regulating the responses to internal and external cues. The transcriptional upregulation of key ABA signaling components, as *ABI4* and *ABI5*, and the corresponding increase in their downstream ABA-responsive genes, such as *EM1*, *EM6*, *RAB18*, and *RD29A*, in the *tcp14 tcp15* mutant further confirms the involvement of TCP14 and TCP15 in attenuating ABA responses. This is reinforced by the fact that a significant enrichment in ABA-responsive, water-deprivation, and stress-related genes was identified among the transcripts upregulated in plants expressing a repressive form of TCP15 (TCP15-EAR), indicating a role for TCP15 in suppressing ABA signaling and stress responses. Interestingly, a large number of ABA-induced genes that contain ABRE sites in their promoters are upregulated in TCP15-EAR plants, suggesting that TCP15 would repress ABA-inducible gene expression through *ABI5* and probably other proteins that bind to ABREs. Furthermore, we found that the introduction of the *abi5* mutation into the *tcp14 tcp15* background significantly reduces ABA hypersensitivity and the expression of the ABI5-target genes *EM1*, *EM6* and *RAB18*, confirming that TCP14 and TCP15 act via *ABI5* to attenuate ABA responses in Arabidopsis.

### 4.2. TCPs repress ABA signaling largely through AFP2

The phenotypes of *tcp14 tcp15* mutant and TCP15 overexpressing seedlings under ABA and salt treatments closely resemble those of mutants and overexpressing lines of negative regulators of ABA responses (Stone *et al*., 2007; Garcia *et al*., 2008; Zhao *et al*., 2021; Li *et al*., 2023; Guo *et al*., 2023; Du *et al*., 2024). For instance, disruption of *AFP2* and *WRKY40*, as well as *RAV1* silencing, results in reduced germination and impaired seedling development under ABA and salt stress, whereas their overexpression confers resistance to ABA (Garcia *et al*., 2008; Shang *et al*., 2010; Feng *et al*., 2014; Lynch *et al*., 2017; Wang *et al*., 2021). Mechanistically, these transcription factors inhibit ABA-dependent responses by directly modulating the expression or activity of ABI transcription factors. Specifically, RAV1 directly suppresses *ABI3*, *ABI4* and *ABI5* expression during seed germination and early seedling growth (Feng *et al*., 2014), while WRKY40 inhibits *ABI5* transcription through the recruitment of the histone demethylase JMJ17 to the *ABI5* locus and by antagonizing HY5-mediated *ABI5* induction (Shang *et al*., 2010; Wang *et al*., 2021). AFP2 has been established as an important negative regulator of ABA signaling that represses ABI5 activity through different mechanisms: by inhibiting its transcriptional expression through changes in histone acetylation (Lynch *et al*., 2017), by promoting its proteasome-mediated degradation (Lopez-Molina *et al*., 2003), and through protein-protein interactions that modulate ABI5 activity (Deng *et al*., 2023). In this study, we revealed that TCP14 and TCP15 repress ABA signaling largely through *AFP2*. This is supported by the fact that *AFP2* expression is downregulated in *tcp14 tcp15* mutants and induced after TCP15 overexpression. Furthermore, AFP2 overexpression largely rescues the ABA-sensitive phenotype in the *tcp14 tcp15* mutant. In addition, our ChIP analyses demonstrated that TCP15 directly associates with a region of the *AFP2* promoter containing TCP boxes. Although AFP2 is a well-established antagonist of ABA responses, its upstream regulators remained poorly established. Our results identify TCP15 as direct upstream activator of *AFP2*, thus adding a new layer to the transcriptional control of ABA signaling. Nevertheless, the partial rescue of *tcp14 tcp15* mutant by AFP2 overexpression indicates that AFP2 is a key, but not exclusive, component of this regulatory module. Indeed, our qPCR and ChIP assays revealed that TCP15 also directly regulates the expression of *RAV1* and *WRKY40*, suggesting that TCP14 and TCP15 activate a broader repression network to modulate ABA signaling. A well-established mechanism of ABI5 action is the positive feedback regulation of its own transcription once activated by SnRK2 kinases, thereby amplifying ABA responses. Thus, it is highly likely that the increased expression of *ABI5* in the *tcp14 tcp15* mutant results from the combined effects of the reduced expression of its negative regulators and the consequent self-activation of *ABI5* expression. Altogether, our findings reveal a transcriptional circuit in which, by directly inducing repressors of ABA signaling, class I TCPs ensure low *ABI5* expression and restrained ABA signaling, thereby promoting optimal seedling growth and developmental progression under non-stress conditions.

### 4.3. Role of TCP transcription factors in abiotic stress responses

Our studies revealed that TCP14 and TCP15 also affect the response to salt-induced abiotic stress conditions. Notably, loss of function of these transcription factors leads to distinct sensitivity to salt depending on the developmental stage. These distinct phenotypic outcomes could be due to the activation of ABA-responsive genes in the *tcp14 tcp15* mutant, which may activate stress adaptation responses but also inhibit growth, especially at early stages of development, as well as seedling establishment. Negative roles in ABA-related responses were previously reported for class I TCPs from several species. For example, Arabidopsis TCP14 interacts with DNA BINDING WITH ONE FINGER 6 (DOF6) to prevent it from activating the ABA biosynthetic gene *ABA DEFICIENT1* (*ABA1*) and other ABA-related stress genes during seed germination (Rueda-Romero *et al*., 2012). Similarly, the class I TCPs StTCP15 and GhTCP19 inhibit ABA biosynthesis to facilitate dormancy release in potato tubers and gladiolus, respectively (Wu *et al*., 2019; Wang *et al*., 2022), while apple MdTCP46 reduces MdABI5 expression and activity, thereby attenuating ABA-mediated drought responses (Liu *et al*., 2022). Nevertheless, a positive role in ABA signaling has also been attributed to other class I TCPs. For instance, OsTCP19 or PeTCP10 overexpression in Arabidopsis enhances ABA sensitivity and drought tolerance (Mukhopadhyay *et al*., 2015; Xu *et al*., 2021), while PCF2 promotes ion homeostasis under stress conditions by upregulating the vacuolar K+-Na+/H+ antiporter NHX1 in rice (Almeida *et al*., 2017). BpTCP20 mediates stomatal closure and ROS scavenging during salt stress conditions in birch (Liu *et al*., 2024). Furthermore, the Arabidopsis class II TCP TCP13 positively regulates dehydration and ABA responses (Urano *et al*., 2022), while BRC1, another Arabidopsis class II TCP, promotes ABA accumulation in axillary buds to prevent branching under limiting light conditions (Gonzalez-Grandio *et al*., 2017). Altogether, these results suggest the existence of a high degree of functional divergence within the TCP family in ABA signaling. Further studies are needed to elucidate the specific roles of different TCP members in ABA signaling and abiotic stress responses, as well as the molecular mechanisms through which they exert their functions in each case.

### 4.4. Functional interplay between ABA and TCP transcription factors

Our results indicate that TCP15, and most likely also TCP14, not only negatively affect ABA responses, but are also targets of ABA-mediated regulation. In fact, we found that ABA negatively affects TCP15 protein levels, but not *TCP15* gene transcription, suggesting that ABA acts post-transcriptionally, possibly by promoting TCP15 degradation. In addition, our expression and reporter assays indicated that TCP14 may be transcriptionally and post-transcriptionally regulated by ABA, assuming that the observed decay in TCP14-GUS levels was significantly more pronounced than the decrease in *TCP14* transcript levels observed in the presence of the hormone. Taken together, our results suggest that ABA negatively affects the abundance of TCP14 and TCP15 proteins, thereby limiting their activity and thus de-repressing the ABA-signaling pathway during times of stress in a double-negative feedback regulatory loop. Furthermore, the ABA-dependent downregulation of TCP14 and TCP15 levels could be a mechanism by which ABA inhibits growth under adverse conditions, since global transcriptomic analysis revealed a group of growth-associated genes that is commonly downregulated by ABA and repressed by a dominant-negative form of TCP15, including known TCP15 targets. This proposal is supported by the fact that ABA barely inhibits the expression of growth-associated TCP-target genes in the *tcp14 tcp15* mutant background, where the expression of these genes is already low under basal conditions, and this is accompanied by reduced ABA inhibition of hypocotyl elongation and cotyledon expansion in the mutant. Therefore, it might be thought that ABA-mediated degradation of TCP proteins could constitute a mechanism to trigger growth arrest under stress conditions, thus ensuring that resources are allocated towards adaptation rather than development. Interestingly, previous studies showed that TCP activity is subject to post-translational regulation in response to environmental and hormonal cues. For instance, TCP15 undergoes redox-dependent inhibition under high-light conditions via oxidation of a cysteine residue in the TCP domain, allowing plants to de-repress the anthocyanin synthesis pathway as a protective response (Viola *et al*., 2013, 2016). Moreover, DA1/DAR peptidases promote TCP14 and TCP15 cleavage to limit cell proliferation, whereas SPINDLY (SPY) stabilizes them to enhance cytokinin responses (Steiner *et al*., 2012, 2016; Peng *et al*., 2015). Conversely, KISS ME DEADLY F-box proteins target TCP14 for degradation in the absence of SPY (Steiner *et al*., 2021). Likewise, DSK2 promotes class I TCPs degradation during plant responses to nitrate starvation (Li *et al*., 2025). These studies suggest that plants fine-tune TCP function at the post-transcriptional level to modulate responses to different environmental and hormonal signals. It would be interesting to elucidate how related are these different reported mechanisms to the ABA-induced degradation of TCPs reported here.

Altogether, our findings support a model of reciprocal antagonism in which TCP14 and TCP15 repress ABA signaling, whereas ABA inhibits TCP-dependent transcriptional responses, including the activation of growth-related genes (Figure 6). This antagonistic interplay between ABA and class I TCPs may be useful to balance growth and stress responses according to environmental conditions. Under favorable conditions, TCP14 and TCP15 promote growth and development by inducing genes involved in these processes, while simultaneously restraining ABA responses by directly inducing repressors of ABA signaling, such as *AFP2*, *RAV1*, and *WRKY40*, which suppress *ABI5* and downstream ABA-responsive gene expression. Under unfavorable conditions, elevated ABA levels lead to reduced accumulation of TCP proteins, relieving ABA signaling repression and thus enabling the activation of ABA-dependent physiological responses, as growth inhibition and stress adaptation. Therefore, by modulating the activity of TCP14 and TCP15, plants can dynamically adjust their responses to changing environmental conditions, prioritizing stress adaptation when needed. This places class I TCPs as crucial players in establishing the balance between growth and stress responses. In a general way, our work contributes to a better understanding of the molecular mechanisms by which plants integrate developmental programs with environmental cues.

**Figure 6.**
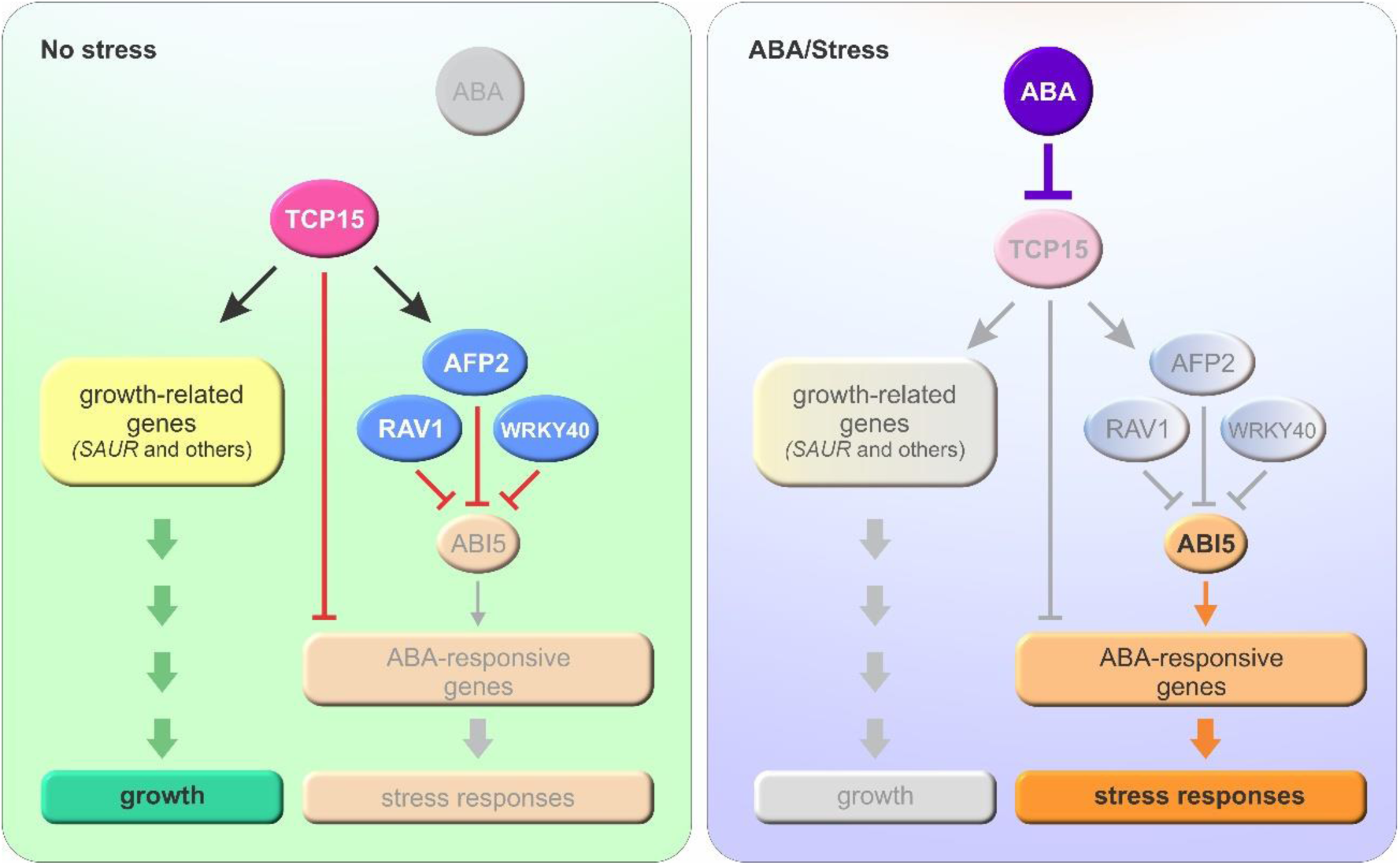
A balance between Class I TCP transcription factors and ABA modulates growth and stress responses. Under non-stress conditions (left), TCP15 (and redundantly TCP14) promotes plant growth by directly activating a group of growth-related genes, while simultaneously inducing the expression of ABA signaling repressors, thereby limiting ABI5 activity and preventing ABA-responsive gene expression. Under rising ABA levels, as those present under certain abiotic stress conditions (right panel), TCP15 protein abundance decreases, thus releasing TCP-dependent repression of the ABA signaling cascade and down-regulating the expression of growth-related genes. This antagonistic relationship provides a mechanism to establish a balance between growth and stress adaptation under changing conditions. Arrow-headed and blunt-ended (T-shaped) lines indicate positive and negative regulation, respectively.

## Acknowledgments

We thank Drs. Martin Kieffer (University of Leeds, UK), Pilar Cubas (Centro Nacional de Biotecnología-CSIC, Universidad Autónoma de Madrid, Spain), Ruth Finkelstein (University of California, USA), and the Arabidopsis Biological Resource Center (Ohio State University), for kindly providing seeds and DNA constructs used in this study.

## Author contributions

ILV and DHG designed the research; AC, JS, MF and ALA performed the experiments; ILV and DHG analyzed the data and wrote the manuscript; all authors read and approved the manuscript.

## Data availability statement

All relevant data are within the article (Figures 1-6) and in the Supporting Information (Figures S1,S2 and Tables S1-S8). This study includes no data that would need to be deposited in external repositories. Sequence data from this article can be found in the Arabidopsis TAIR database under the accession numbers: *TCP14*, AT3G47620; *TCP15*, AT1G69690; *ABI4*, AT2G40220; *ABI5*, AT2G36270; *EM1*, AT3G51810; *EM6*, AT2G40170; *RAB18*, AT5G66400; *RD29B*, AT5G52300; *AFP2*, AT1G13740; *WRKY40*, AT1G80840; *RAV1*, AT1G13260; *SAUR14*, AT4G38840; *SAUR50*, AT4G34760; *SAUR51*, AT1G75580; *SAUR63*, AT1G29440.

## Supporting Information

**Figure S1.** ABA sensitivity of single mutants in *TCP14* and *TCP15*.

**Figure S2.** GUS histochemical analysis of TCP14 and TCP15 expression under NaCl treatment.

**Table S1.** Oligonucleotides used in this study.

**Table S2.** Gene Ontology (GO) enrichment analysis on differentially expressed genes in *pTCP15::TCP15-EAR* plants.

**Table S3.** List of "Response to abscisic acid" differentially expressed genes in *pTCP15::TCP15-EAR* plants.

**Table S4.** List of "Response to water deprivation" differentially expressed genes in *pTCP15::TCP15-EAR* plants.

**Table S5.** List of "Response to stress" differentially expressed genes in *pTCP15::TCP15-EAR* plants.

**Table S6.** Gene Ontology (GO) enrichment analysis on commonly upregulated genes in *pTCP15::TCP15-EAR* plants and ABA-treated wild-type plants

**Table S7.** Gene Ontology (GO) enrichment analysis on commonly downregulated genes in *pTCP15::TCP15-EAR* plants and ABA-treated wild-type plants

**Table S8.** List of commonly downregulated genes related to growth in *pTCP15::TCP15-EAR* plants and ABA-treated wild-type plants

