## Supplementary Figures for "Reciprocal regulatory interactions between class I TCP transcription factors and ABA signaling balance growth and stress responses in Arabidopsis"

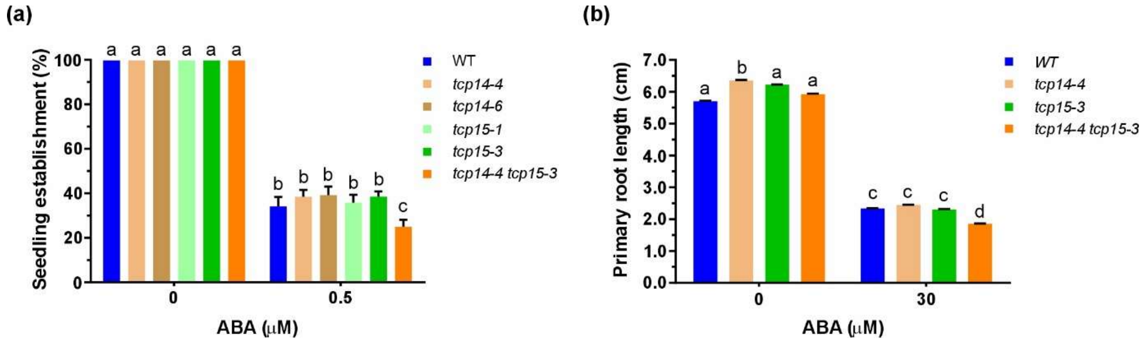


**Figure S1.** ABA sensitivity of single mutants in *TCP14* and *TCP15*. (A) Effect of ABA on seedling establishment in plants of the indicated genotypes grown for 10 d in the absence or presence of 0.5μM ABA. (B) Effect of ABA on primary root elongation. Seedlings grown during 3 d in the absence of ABA were transferred to MS medium supplemented with the indicated concentrations of ABA. Root length was measured 5 d after transfer. Bars show the mean±SD. Different letters denote statistically significant differences ($P < 0.05$; ANOVA). All the experiments were repeated three times with similar results.

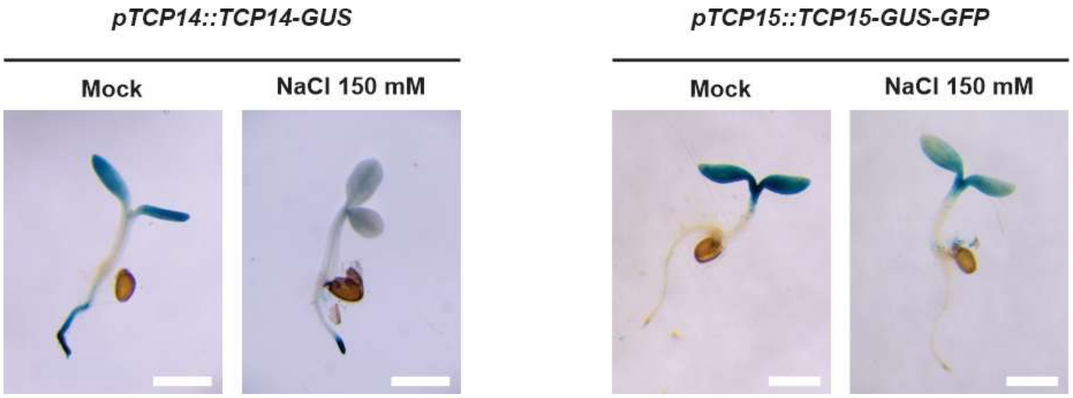


**Figure S2**. GUS histochemical analysis of TCP14 and TCP15 expression under NaCl treatment. TCP14 and TCP15 expression patterns were analyzed by β-glucuronidase (GUS) histochemical staining of *pTCP14::TCP14-GUS* and *pTCP15::TCP15-GUS-GFP* seedlings grown during 6 d in liquid MS medium under normal conditions and then treated with either 150 mM NaCl or H2O (Mock) for 24 h. The images are representative of 10 seedlings analyzed (scale bars: 1 mm). The experiment was repeated three times with similar results.
